# Characterization of a novel amastin-like surface protein (ALSP) of *Leishmania donovani*, a probable lipase

**DOI:** 10.1101/2020.07.23.218107

**Authors:** Bapi Biswas, Bhakti Laha, Pratyay Sengupta, Koushik Das, Monidipa Ghosh

## Abstract

In the current study, a novel putative protein of *Leishmania donovani*, amastin-like surface protein (ALSP) has been characterized. The gene was cloned in a bacterial system and the protein was overexpressed. A polyclonal antibody was developed against the protein, which detected a 10 kDa band in the *L. donovani* amastigote. ALSP mRNA was detected in *L. donovani* amastigote, which was not expressed in the promastigote. ALSP mRNA was not expressed in either morphological forms of *Leishmania major*. MALDI-TOF confirmed the molecular weight of ALSP as 10 kDa. I-TASSER predicted the function of ALSP as a lipase, which was confirmed by preliminary *in-vitro* experiments using amastigotes of *L. donovani*. ALSP has GAS amino acid sequences, which might act as the active site for its lipase activity. The selective expression of ALSP in amastigotes probably makes it important in virulence mechanisms such as survival in the phagolysosome and modulation of its membrane and other metabolic functions, necessary for parasite survival in the human host. ALSP can act as a peptide vaccine target and maybe detected in the peripheral blood or urine as a molecular biomarker of active disease in visceral leishmaniasis.

## INTRODUCTION

Leishmaniasis is an insect-borne global scourge, caused by the protozoan parasite *Leishmania*. The morphology of different species of *Leishmania* alternates between two distinct morphs: an extracellularly located flagellated promastigote, swarming in the gut of female phlebotomine sandflies and morphologically distinct intracellular located non-flagellated amastigote replicating in the macrophages of the mammalian host ^[1][2]^. World Health Organization (WHO) reports that 12 million individuals are affected by leishmaniasis around the globe, among which 20,000–40,000 people are decimated by the disease every year^[3]^. Limited drugs such as antimonials, amphotericin B, and miltefosine are available to treat visceral leishmaniasis (kala-azar) but they provide limited efficacy due to toxic effects, nonspecific modes of action, increasing drug resistance, and rising cost^[4]^. Till date no vaccine has been developed against the disease ^[5] [6]^.

Kinetoplastids like different species of *Leishmania* and *Trypanosoma* express a wide family of surface glycoprotein called amastin. *In-silico* prediction studies have revealed a low molecular weight analog called amastin-like surface protein (ALSP) (Gene ID: LDBPK_34265)^[8] [9]^. Whether at all ALSP is related to amastin is hitherto unknown. Previous studies have shown that the immunogenicity of amastin sequences are the furthermost among all surface antigens of *Leishmania* in mice^[9]^ and show intense immunogenicity with human visceral leishmaniasis^[10]^. This low molecular weight protein may be an efficient target for vaccine candidacy^[11]^. Diverse amastin protein are detected in *L. donovani* amastigotes in visceral leishmaniasis patient ^[12] [13]^. Our preliminary studies on structural prediction highlighted a lipase-like role for ALSP. Some previous studies have identified lipases as a key virulence factor for the live amastigote in the host phagolysosome and coordination of complex metabolism.

The lipid turnover helps the macrophage to eliminate the intracellular amastigote. Constituent lipids of surface membranes (SM) of the parasite have unique properties contributing to the organism’s resistance to digestion in the hydrolytic environs of both its insect vector and mammalian hosts ^[14] [15] [16]^. Lipases may also modulate membrane properties and tissue modelling to favour widespread dissemination, facilitating in the visceralization of the parasite^[17][18] [19]^.

In the current study, we characterized the novel amastin-like surface protein (ALSP), expressed its gene in a bacterial system, overexpressing the protein *in-vitro*, determined its molecular weight and expression patterns in *L. donovani* promastigotes and amastigotes evaluated and its predicted function as a lipase.

## MATERIALS AND METHODS

### Culture of cells and parasites

Promastigotes of *L. donovani* (AG83) were maintained at 23 °C in Medium199 (M199), with 10% FBS (Fetal Bovine Serum), penicillin (50 U/mL), and streptomycin (50 μg/mL) (Gibco, US). Centrifugation at 1000*g* was done for 10 min to obtain promastigote pellets in their late log phase (10 million promastigotes/mL). Then, it was washed with phosphate buffer saline (PBS) at pH 7. The promastigotes of *L. major* was cultivated in M199 media plus 4-(2-hydroxyethyl)-1-piperazineethanesulfonic acid (HEPES) buffer at 25 °C. Human acute monocytic leukemia cells THP1 was maintained in RPMI (Gibco, US) media, with added FBS and the antibiotics like penicillin and streptomycin. *In-vitro* conditions changed the promastigotes into amastigotes within the THP1 cells at a ratio of 1:10 (cell: parasite). The amastigotes were isolated by the procedure, optimized in the laboratory, following the methods described by Moreno MLV et al. with some modifications^[20]^.

### Accession numbers

The IDs (Identities) and annotations of the novel protein sequences are as follows: GeneDB^i^(CBZ37742.1, LDBPK_342650, XP_003864424.1), Gene ID: 13392833

### Prediction of ALSP 3D structure and function through iterative threading assembly refinement algorithm (I-TASSER)

ALSP amino acid sequence was uploaded to the I-TASSER server (https://zhanglab.ccmb.med.umich.edu/I-TASSER/)^[21]^. A large ensemble of structural conformations, called decoys is generated by I-TASSER stimulations for each target. The SPICKER program uses the decoys to select the best five final models based on pair-wise structure similarity. The confidence score (C-score) is used to determine the structure quality, which varies between −5 to 2. Higher C-score defines a model as more significant than a lower one. Template modelling score (TM-score) and Root Mean Square Deviation (RMSD) are estimated by confidence score (C-score) and protein length following the correlation between these parameters.

Based on the I-TASSER structure prediction the function of the ALSP has been annotated by COFACTOR and COACH. COFACTOR infers protein functions (ligand-binding sites, EC and GO) using structure comparison and protein-protein networks. COACH uses a meta-server approach that combines multiple function annotation results from the COFACTOR, TM-SITE, and S-SITE programs.

### Model evaluation

The predicted 3D structure of ALSP was evaluated by PROCHECK (https://www.ebi.ac.uk/thornton-srv/software/PROCHECK) and ProQ (https://proq.bioinfo.se/cgi-bin/ProQ/ProQ.cgi) web servers. PROCHECK is used to test the stereochemical quality, correctness of the 3D protein structure ^[22]^, and the validation of generated models was further accomplished by ProQ ^[23]^.

### Lipase assay

Triglyceride lipase is an enzyme that hydrolyses triglycerides into glycerol and fatty acid ^[24]^. The enzyme activity of purified ALSP from *L. donovani* amastigotes in whole lysate was checked by Lipase Assay Kit (R_EAl_G_ENE_, US, Cat # 250069) expressed in Unit/μg. The Unit/μg (U/μg) is defined as 1μg protein-producing 1umol glycerol/fatty acid in 1min at 37 °C. The ALSP was purified from the whole cell lysate of the *L. donovani* amastigotes through immunoprecipitation assay which followed the same protocol described below for affinity-based immune-precipitation assay for the native protein purification. The whole-cell lysate of *L. donovani* promastigotes was taken as the negative control in the respective experiments

The experiment was performed with blank control (B), test tube (T), and standard tube (S). The OD_710_ values of blank control were subtracted from the standard tube OD_710_ and test tube OD_710_ values.

The 125 μM/mL of oleic acid standard solution was diluted to 62.5, 31.25, 15.625, 7.8125, and 3.9 μM/mL with anhydrous ethanol to make a standard curve at 710 nm absorbance. The standard curve was used to affirm the enzyme activity of the purified ALSP in amastigote form.

### Determination of Antigen efficiency as Vaccine Candidate

The antigenicity of ALSP was checked through ANTIGENPro and VaxiJen 2.0 online tools^[25] [26]^

### Cloning the ALSP gene

The ALSP gene (Gene ID: CBZ37742.1) was PCR (polymerase chain reaction) amplified in 50 uL reaction volume containing 100 ng of purified genomic DNA of *L. donovani* promastigotes. 100 pmol forward (EcoRI->5’GT C GAATT CGTATG CAT ATG CGT GTA CTT GTG CGT 3’) and reverse primer (XhoI->5’TTCTCGAG TCA GCA CGG AAA GGA ACG CGA 3’), 200 uM dNTP (Deoxynucleotide triphosphates), 3U Taq DNA polymerase. The fragment was incorporated into the pGEX4T2 vector at its multiple cloning site, using the restriction sites of EcoRI and XhoI. The pGEX4T2-ALSP hybrid construct was used to transform the *Escherichia coli* DH5α and BL21 respectively.

### Confirmatory analysis of Cloning through colony PCR, double digestion of the cloned Plasmid, and DNA sequencing

ALSP-forward and reverse primers were used to do colony PCR with transformed and untransformed DH5α and BL21 cells, following the same PCR program. The restriction-digestion of the ALSP-GST construct was done after isolating the plasmid from the transformed DH5α cells, using a plasmid isolation mini kit (GCC Biotech). The isolated plasmid was dissolved in 20 μl of Diethyl pyrocarbonate (DEPC)-treated water and used for Sanger sequencing (Xcelris Labs, Bangalore, India). Nucleotide-nucleotide basic local alignment search tool (BLASTn) was performed for the chromatogram to check the identity with the available sequence of *Leishmania* species in the National Center for Biotechnology Information (NCBI) database.

### Production of anti-ALSP antibody and substantiating analysis of its specificity

Antisera were developed against the ALSP by an optimal peptide epitope domain of a protein (SSPFSSTRSSSSSRS –C), the cysteine residue addition at the C terminal end is required for keyhole limpet hemocyanin (KLH) conjugation, applied by a bioinformatics tool (BioBharati LifeScience Pvt. Ltd. Kolkata, India). The conjugation was used to immunize New zealand White Rabbit. Preimmuned and immunied serum was realized up to 6^th^ booster dosage (till 2 and half months). Antibody titers were evaluated by indirect ELISA. 500 pg of ALSP per well was applied to study their titer of different dilutions factor (1:500, 1:1000, 1:2000, 1:5000, 1:10000, 1:20000, 1:400000 dilutions) of antisera containing antibody of both the batches.

### Affinity-based immune-precipitation assay for the native protein purification

1×10^6^ cells of promastigote and amastigote of *L. donovani* were resuspended in 1x PBS (PH-7.4) and kept at 4 °C for 10min. Parasites were lysed through repeated freeze thawing. Supernatants were collected by centrifugation at 100 g and 200 g respectively for amastigotes and promastigotes. Then the collected supernatants were used for immunoprecipitation with the antisera (ALSP) raised in of New Zealand White rabbit and antisera contains albumin-like protein. The immunoprecipitation was carried out with Protein A-Sepharose bead, added to both the antisera in a 1:2 volume ratio and incubated at 4 °C overnight with gentle inversion rotation. Unbounded antisera were washed out by using 1xPBS (PH-7.4). This process was repeated thrice at room temperature and then centrifuged at 200 g for promastigote and incase of amastigote centrifuged at 100 g. The soluble supernatants of both forms were added to the specific antibody-loaded Protein A-sepharose column and incubated overnight at 4 °C with gentle inversion rotation. The unbound protein was removed. The bound protein with antisera through Protein A-sepharose bead was separated by boiling with sodium-dodecyl-sulfate-polyacrylamide gel electrophoresis (SDS-PAGE) loading buffer for 15 min at 100 °C and collected by centrifugation. The elutes were analyzed by immunoblotting after incubation with the anti-ALSP polyclonal antibody raised against ALSP. The albumin-like protein was used as the negative control as it is absent in *L. donovani* amastigotes ^[27]^.

### MALDI-TOF (Matrix-Assisted Laser Desorption/Ionization-Time of Flight)

The protein dialysate was subjected to MALDI-TOF to detect the molecular weight. The solubilized protein was added to a target MALDI plate matrix using α-cyano-hydroxycinnamic acid (CHCA). This was examined by MALDI-TOF mass spectrometer (Applied Biosystems, USA). The resultant spectra were visualized. The study was done at the central instrumentation facility at the Council of Scientific & Industrial Research-Indian Institute of Chemical Biology (CSIR-IICB) Kolkata.

### Determination of native protein through sequencing

The MALDI-TOF-MS/MS analysis was done using trypsin digested native ALSP sample to check its sequence identity based on MS/MS Ions Search on the MASCOT database server (http://www.matrixscience.com/search_form_select.html). Parameters considered were: carbamidomethyl (C) for fixed modification, oxidation (M) for variable modification, trypsin as enzyme, peptide mass tolerance (100 ppm), and fragment mass tolerance (+0.2). This study was carried out at CSIR-IICB Kolkata using the central instrumentation facility for mass spectrometry (MALDI-TOF-MS/MS).

### Two-step reverse transcription PCR

ALSP transcripts in both forms the of *L. donovani* were examined using TRI Reagent (Sigma T9424) isolated whole-cell RNA. RNA was reversed transcribed to cDNA at 42 °C for 60 min. Forward and reverse ALSP gene-specific primers were used. Polymerase Chain Reaction (PCR) amplification was attempted in the next step.

### Fluorescence microscopy

Digenic forms of *L. donovani and L. major* were obtained by centrifugation (4000 g for promastigotes and 100 g for amastigotes). The pellets were washed twice with phosphate buffer saline. After fixation with chilled methanol for 2 min, the cells were treated with permeabilization buffer for 30 sec to 1 min. The pellets were rewashed with phosphate buffer saline and conjugated with preimmune sera and immune sera, targeted against ALSP (1:25 dilution) in the presence of blocking agent 3% bovine serum albumin for 45 min. The cells were assessed with fluorescein isothiocyanate (FITC)-conjugated goat-derived IgG (Thermo Fisher Scientific, Waltham) at 1:500 dilution for 30 min. Cells were viewed after cover slipping with mounting media plus DAPI and viewed with a fluorescence microscope (Zeiss, UK) at 40X. Preimmune sera was used as negative controls. Appropriate optical filters were used for 4’,6-diamidino-2-phenylindole (DAPI) (λ = 461 nm) and Fluorescein isothiocyanate (FITC) (λ = 591 nm) respectively.

## RESULTS

### Prediction of 3D structure and function of ALSP and evaluation of the Model by PROCHECK and ProQ

**Figure 1.a** showed the Helix (H), Coil (C), and Strand (S) regions of the predicted secondary structure of ALSP and their solvent accessibility. The 3D structure of the ALSP protein was projected by homology modelling using its amino acid sequence on the I-TASSER. The range of the C-score is −5 to 2, where a model is highly significant with a magnificent value of c and conversely. C-score of top 5 models were −3.58, −4.36, −4.77, −4.56, and −4.95 respectively. Model 1 contains a higher C-score, as shown here (**Figure 1.b**). The function of ALSP was determined *in-silico* by COFACTOR and COACH software, derived from the I-TASSER projection. The likely function of ALSP was triglyceride lipase, (GO: 0004806). The biological process is associated with cytosolic lipid metabolism (GO: 0016042). The lipase enzyme active site of the ALSP was predicted to be the GAS amino acid’ sequence (**Figure 1.c**). A Confirmatory study of the ALSP 3D structure was done by the construction of Ramachandran plots through PROCHECK. The result revealed that residues of the ALSP model fall within 79.1% and 19.4% in the most favored and additional allowed regions independently, and there was only 1.5% of residues that fall within disallowed regions (**Figure 1.d**). In broad-spectrum, a score of about 100% indicates the best stereochemical quality of the model ^[28]^. Consequently, the PROCHECK result proposes that the predicted model quality of ALSP is satisfactory. The predicted 3D structure quality was also checked via ProQ and it is evident that the predicted LG score was 2.193, which is greater than 1.5. The ProQ database confers a fairly good model with a more than 1.5 LG score. The antigenicity of ALSP was 0.7883 and 0.6494, calculated by using VaxJen and AntigenPro software.

**Figure. 1.**
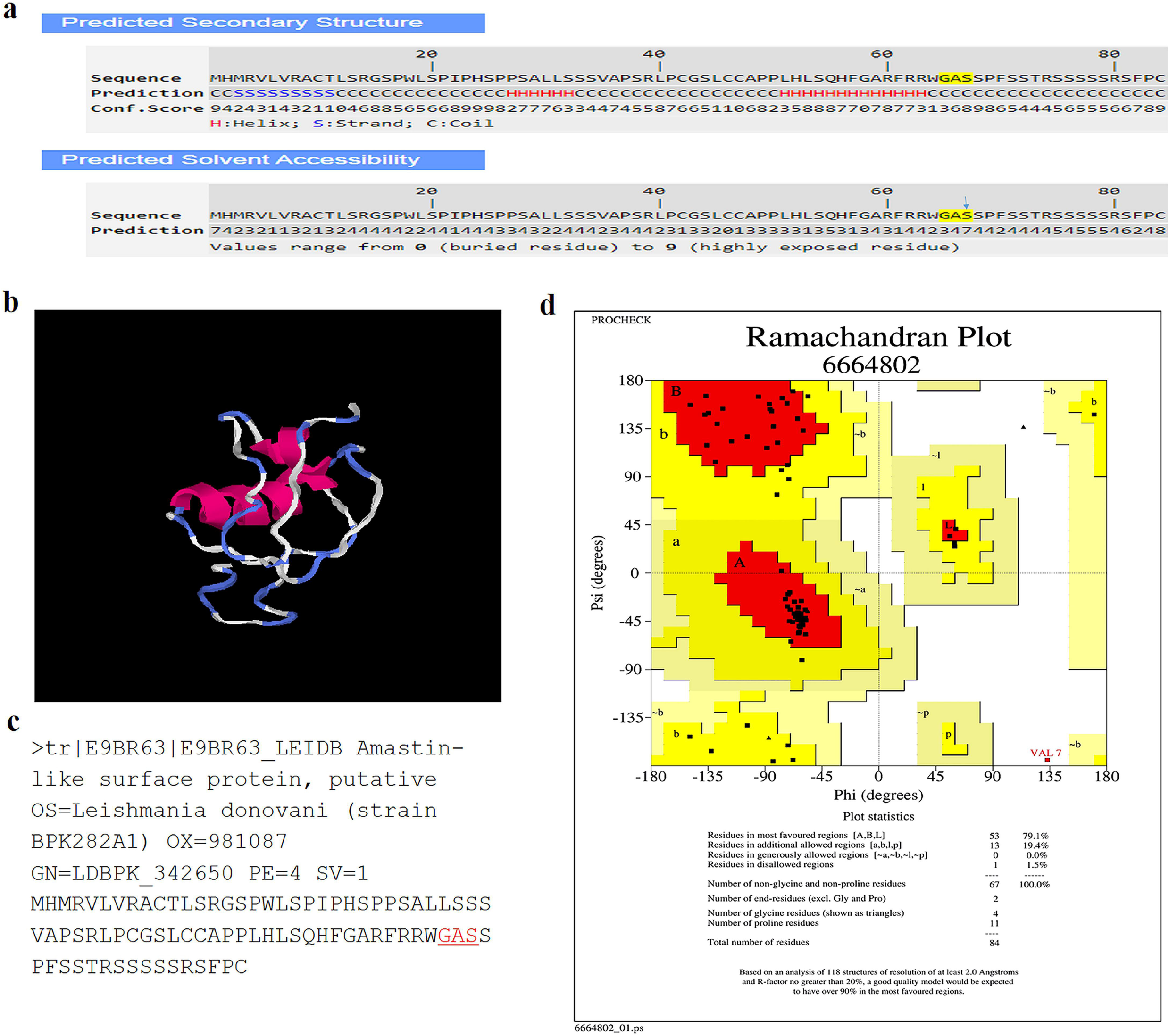
Structure prediction and structure-based function annotation of ALSP through I-TASSER. (1.a) Predicted secondary structure and solvent accessibility of ALSP (1.b) Predicted best model of ALSP by I-TASSER where C-score is −3.58, estimated TM-score =0.32Â±0.11, and estimated RMSD-score=11.6Â±4.5Ã. (1.c) Exhibited GAS region of ALSP from its amino acids sequence (1.d) Validation of ALSP predicted 3D model through Ramachandran plot where 79.1% of ALSP residues fall within the most favoured and 19.4% in the additional allowed regions independently, and only 1.5 % of residues that fall within disallowed regions.

### Pilot study of In-vitro functional analysis of ALSP through Lipase assay

The amino acid sequence of ALSP was analyzed with the I-TASSER software for predicting its probable role (**Figure 2.a**). Based on the *in-silico* data, the function of ALSP was examined through *in-vitro* analysis by Lipase assay (**Figure 2.b**). Oleic Acid was used as a standard solution. The enzyme activity of purified ALSP in *L. donovani* amastigotes was examined through the standard curve made by different concentrations of oleic acid at OD_710_. The crude cell lysate of *L. donovani* promastigotes was used as negative control. The purified ALSP produced fatty acid in *L. donovani* amastigote, while the crude promastigotes almost did not show any such activity for ALSP. 0.98 mg/mL, 1 mg/mL, and 0.87 mg/mL amastigotes proteins were purified through immunoprecipitation assay using raised polyclonal antibody against ALSP, produced 13.59 μmol/mL, 11.1676 μmol/mL, and 11.32 μmol/mL fatty acid. Subsequently, the enzyme activity was calculated for ALSP in the amastigote form, and it was estimated to be 1.3869 U/mg, 1.1167 U/mg, and 1.3018 U/mg. The mean enzyme activity was 1.2684 U/mg for purified ALSP in *L. donovani* amastigotes and crude lysate in *L. donovani* promastigotes was shown 0.0000048 U/mg **(Figure 2.c**). The standard deviation in the sample was 0.130249.

**Figure. 2.**
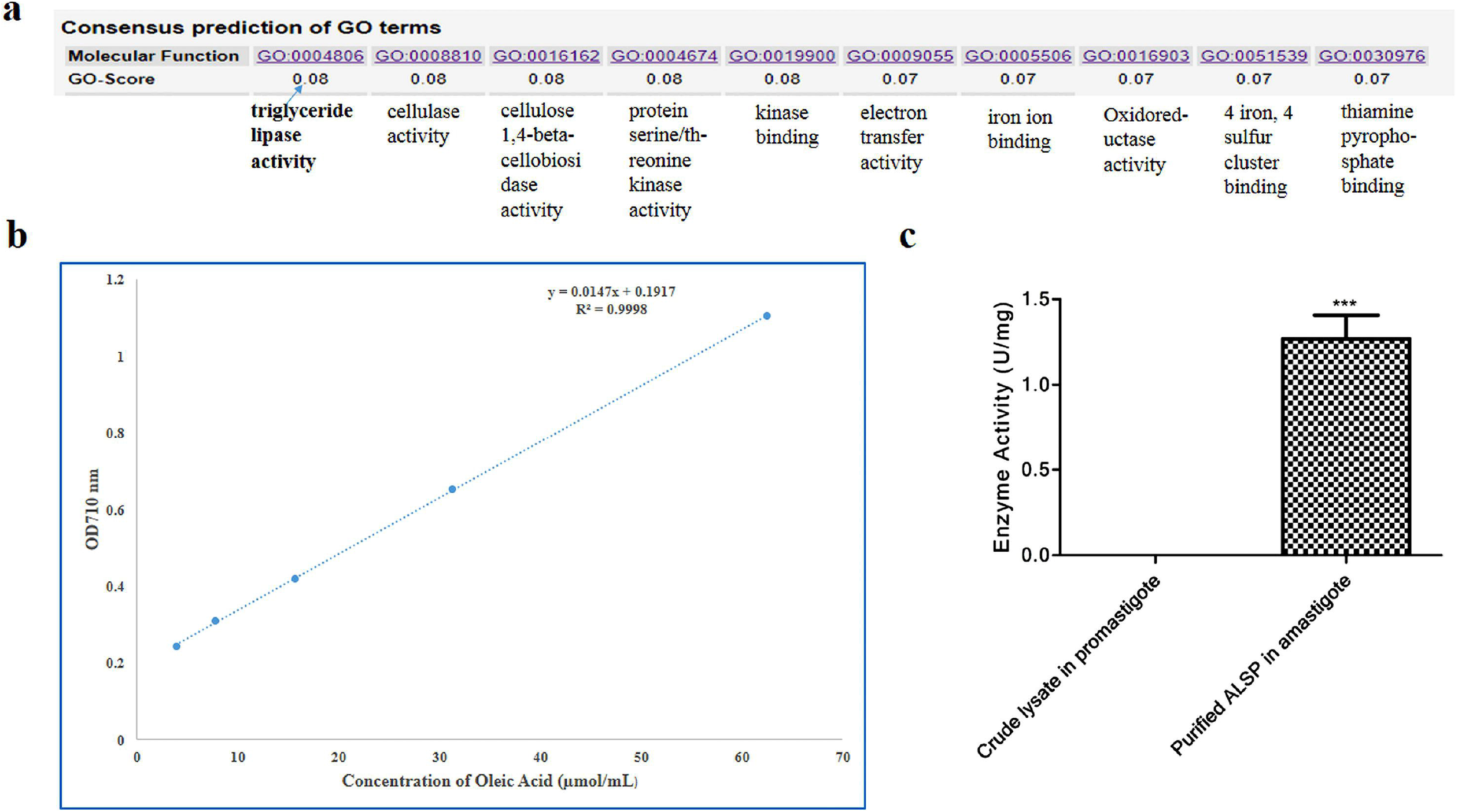
Function of ALSP as triglyceride lipase. (2.a) The function of ALSP predicted through *in-silico* analysis using its amino acids sequence in I-TASSER online tool. It was found to act as a lipase enzyme with other possible roles. Fig. 2.b Depicts a standard curve between different concentration of Oleic Acid and Absorbance of Oleic Acid at 710nm. (2.c) Lipase activity of purified ALSP in amastigote and promastigote forms of *L. donovani*. It showed 1.3869 U/mg lipase activity of purified ALSP of amastigotes. But promastigotes specific protein did not show such kind of activity. The standard deviation in the sample was 0.130249. The Two-tailed T-test exhibited the p-value 0.0001 for the purified ALSP in amastigote form and which is highly significant.

### Overexpression of the leishmanial complete gene (CBZ37742.1) of ALSP and confirmation

The ligated product of the pGEX-4T-2 vector and ALSP gene was transferred on LB-AMP plate with DH5α competent cells (**Figure 3.a**). Thereafter, the cloning of the ALSP gene was confirmed through colony PCR, restriction digestion, and sequence analysis. The colony PCR with colony 1 and colony 2 visualized a sharp band for each near 255 bp (**Figure 3.b**). Consequently, those two colonies were used for restriction digestion with EcoRI and XhoI restriction enzymes, and the restriction digestion result was exhibited an intense band close by 255 bp and 4970 bp for the ALSP gene and pGEX-4T2 vector in respect of 1 kb DNA Marker. The chromatogram confirmed the cloning of ALSP in the pGEX-4T2 vector with 89.12% identity (**Figure 3.c & 3.d**) in respect to the *L. donovani* database. The target gene was expressed in the *E. coli* BL21 strain and used for protein purification in abundant quantity.

**Figure. 3.**
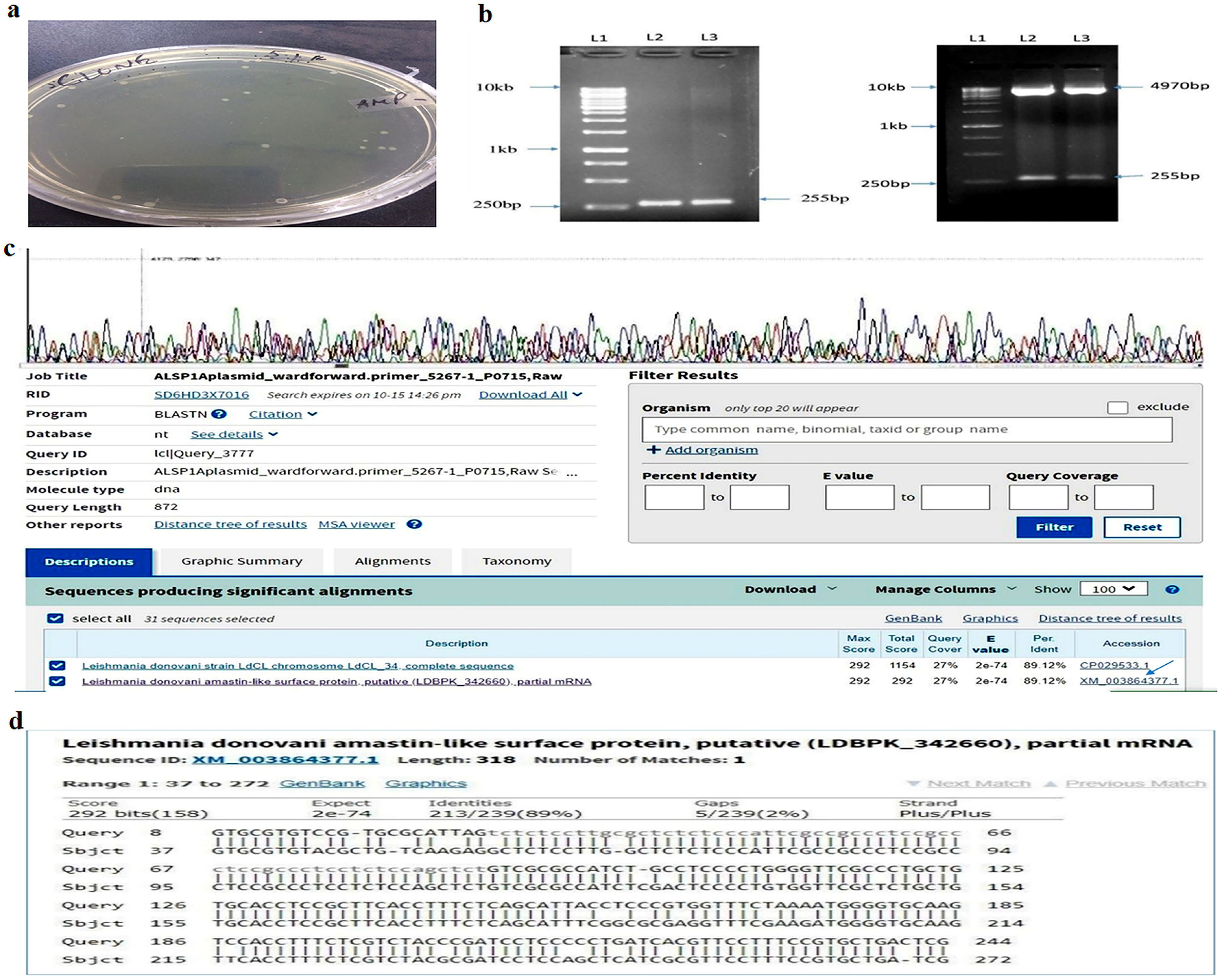
Confirmation of the cloning in a prokaryotic system with transformed DH5α competent cells through colony PCR, restriction digestion, and sequencing analysis. (3.a) Transferred ligated pGEX-4T-2 vector and Gene of Interest (ALSP) on LB-AMP plate with *E. coli* DH5α competent cells (3.b) Colony PCR of cloned plasmids of colony 1 (Lane 2) and colony 2 (Lane 3) and diagram of restriction digestion of those cloned plasmids with EcoRI and XhoI restriction enzymes exhibited an intense band near 255 bp for each colony in respect of the 1 kb DNA ladder at Lane 1. (3.c and 3.d) Chromatogram of pGEX4T2 vector with the 255 bp long ALSP insert and Sequencing analysis of the ALSP gene through Nucleotide BLAST against Leishmanial gene data bank.

### Production of antiserum against epitopic ALSP sequence

The highly probable antigen of KLH conjugated ALSP was used for immunization of NZ White Rabbit Model. Antibody concentrations were calculated. Affinity purification of antisera was performed by indirect ELISA. Both batches demonstrated a diminishing titer upon and increasing dilution. The concentration of preimmune sera remained insignificant in comparison with the immunized batches of animals, which implied that the developed antisera was complementary against the antigen. The specificity was checked using Indirect ELISA (**Figure 4.a and 4.b**). Batch 1 showed a higher titer than Batch 2 (**Figure 4.a**). Batch 1 was used to purify ALSP-antibody through true affinity purification (**Figure 4.b**).

**Figure 4.**
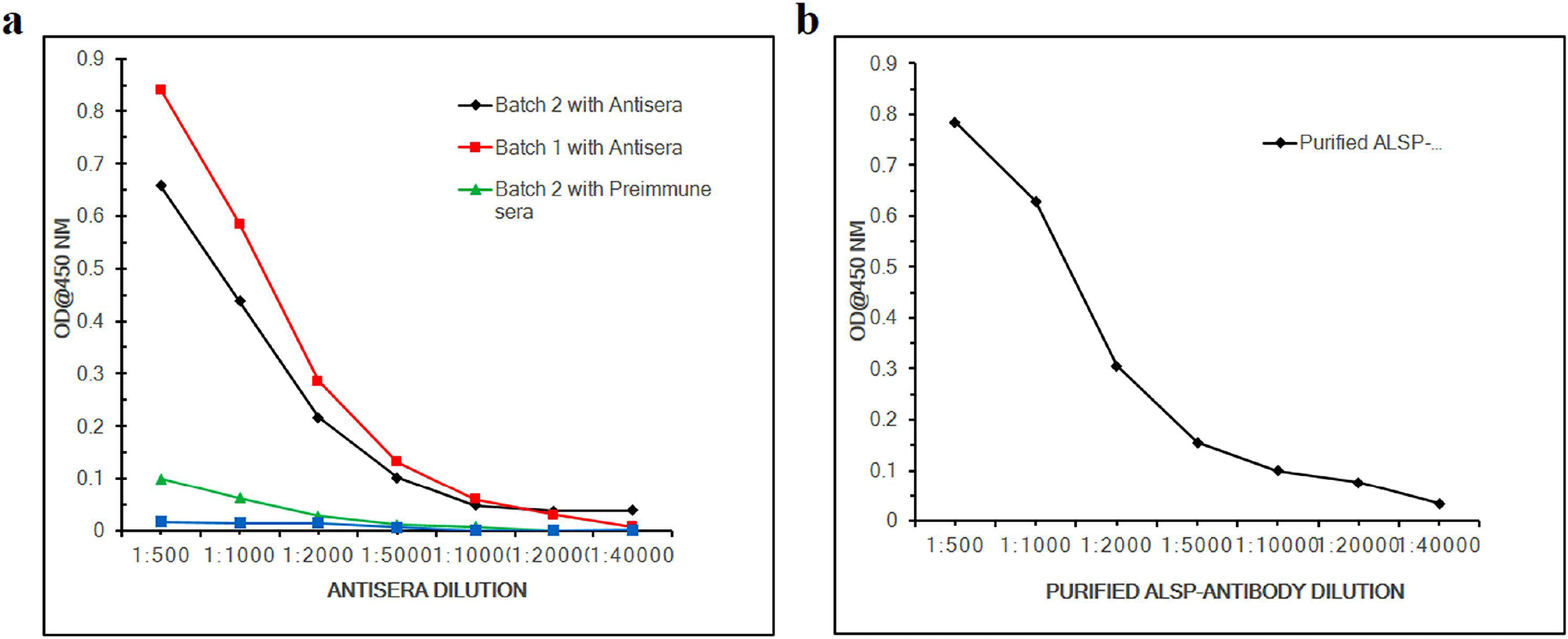
Evaluation of polyclonal antibody against Leishmania donovani ALSP protein. (4.a) Binding dynamics of polyclonal antibodies evaluated by indirect ELISA. The titer of preimmune serum is highlighted by the green and blue traces. The binding strengths of antisera in batch 1 and batch 2 are projected by red and black traces respectively. (4.b) Affinity purification of ALSP-antibody using antisera from batch 1.

### Determining ALSP gene and protein expression in *Leishmania donovani*

The gene transcription of ALSP in the promastigote and amastigote of *L. donovani* were analyzed by reverse transcription PCR. GAPDH gene (496bp) was used as a positive control for the promastigote and amastigote of *L. donovani* respectively (**Figure 5.a.II**). Reverse transcription PCR product of the ALSP gene using complementary DNA (cDNA) from amastigote form of *L. donovani* revealed a band at 255 bp (**Figure 5.a, L3**). No band was seen for the lysate derived from the promastigotes (**Figure 5.a, L2**). This demonstrated that the ALSP mRNA expression was exclusively in the amastigotes.

**Figure. 5.**
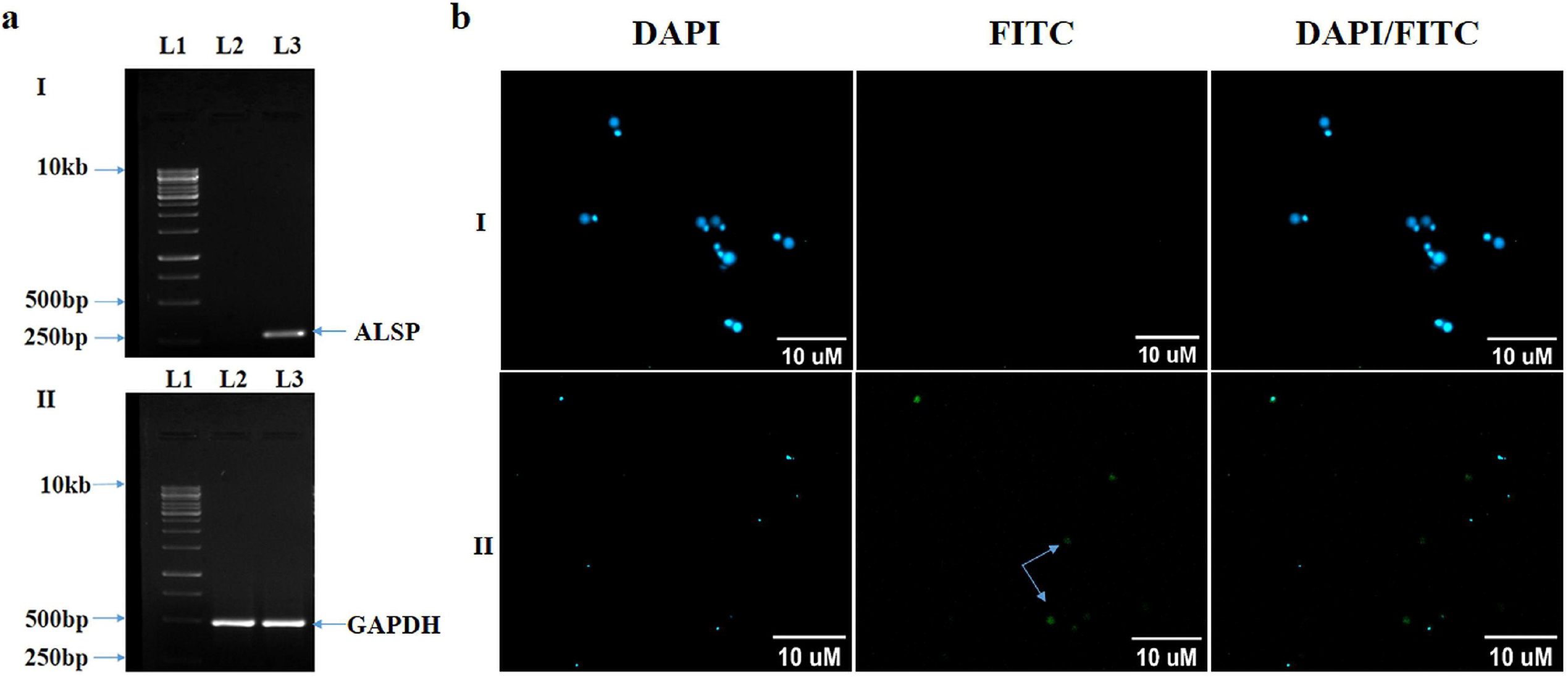
Determining ALSP gene and protein expression in Leishmania donovani. 5.a.I. Reverse transcription-based amplification of ALSP-encoding gene transcripts using whole-cell RNA obtained from promastigote and amastigote forms of *L. donovani*. Lane 1, 1 kb DNA marker; Lane 2 & 3, RT-PCR products of whole-cell RNA using cDNA from promastigotes and amastigotes respectively. Lane 3 contains a sharp band near 255 bp; no band detection in Lane 2; 5.a.II. Illustration of positive control for both structural forms of *L. donovani*. Lane 2 & 3 RT-PCR products of whole-cell RNA of GAPDH in promastigote and amastigote forms respectively, Lane 1, 1 kb DNA marker. 5.b. Fluorescence microscopy also shows the same result while the native protein is probed with antisera and FITC conjugated goat-derived anti-rabbit IgG secondary antibody using the filters at 461 nm for DAPI and 591 nm for FITC. 5.b.I. Represents promastigotes with antisera whereas 5.b.II. Exemplifies amastigotes with antisera. The arrows indicate the expression of ALSP in amastigotes.

The expression of ALSP in promastigote and amastigote forms was examined through fluorescence microscopy with antisera and FITC-conjugated anti-rabbit IgG secondary antibody at 40x magnification with mounting media containing DAPI. It revealed the expression of ALSP protein in amastigotes (**Fig. 5.b.II**) but not in promastigotes. (**Figure 5.b. I**).

### Determining ALSP gene and protein expression in *L. major*

The transcript level expression of the ALSP gene was checked in both structural forms of *L. major* to confirm its specificity. It was done through reverse transcription PCR (RT-PCR), using cDNA acquired from the RNA of both the vertebrate and invertebrate forms of *L. major*, considering GAPDH (496 bp) as loading control (**Fig. 6.a.II**). The Reverse transcription PCR products of ALSP with the gene-specific primers were run on a 1% agarose gel, which exhibited no band for amastigote and promastigote (**Figure 6.a**, **L5 & L4**) of *L. major*, but the amastigote of *L. donovani* showed a clear band nearby 255 bp (**Figure 6.a.I**, **L3**) for ALSP.

**Figure. 6.**
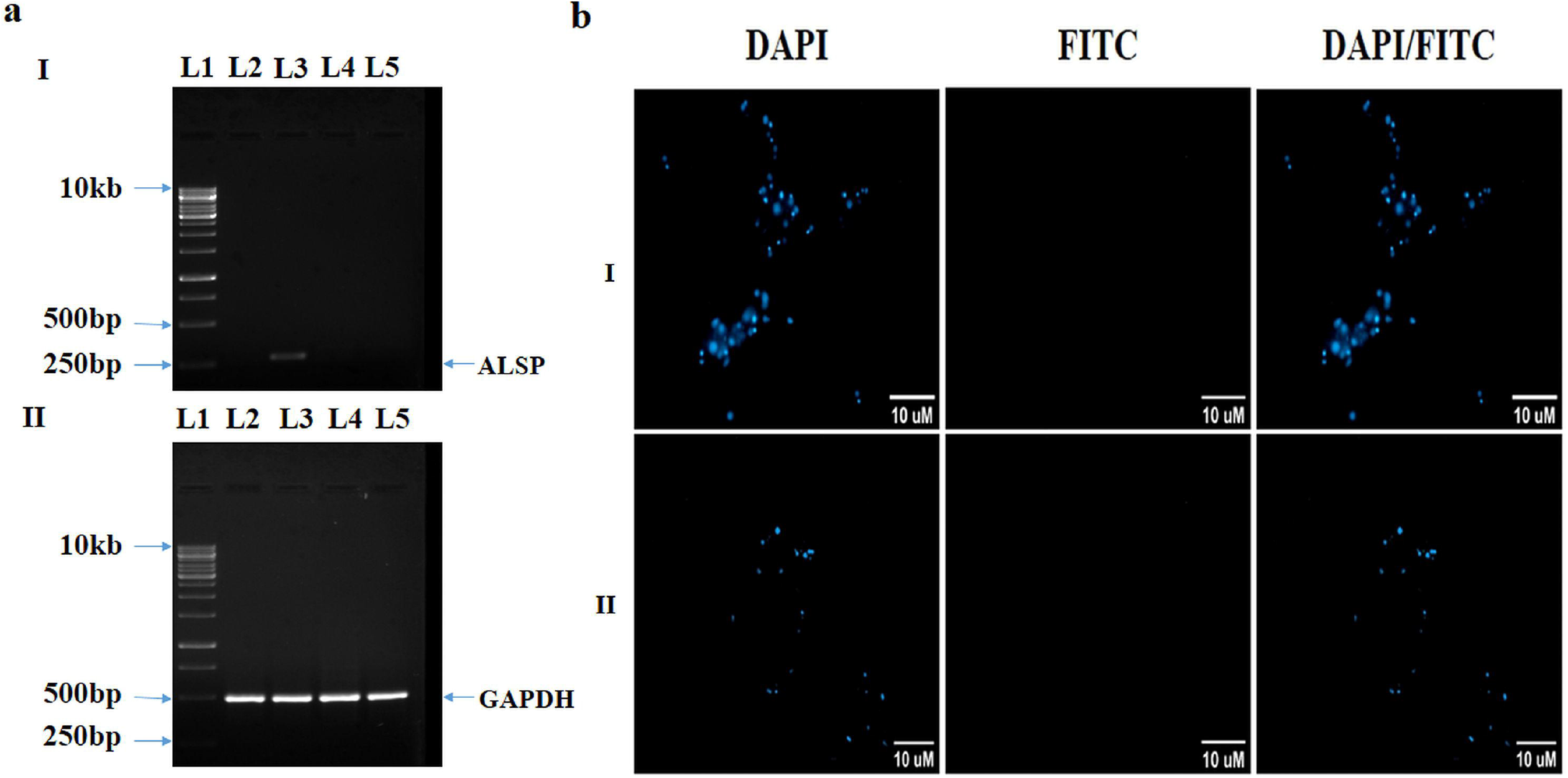
Determining ALSP gene and protein expression in Leishmania major. 6.a.I. RT-based enhancement of ALSP transcripts by total RNA acquired from both morphological forms of the parasite *L. major*. Lane 1, 1 kb DNA marker; Lane 2 & 3, RT-PCR products of whole-cell RNA using cDNA from both promastigote and amastigote structural forms of *L. donovani* respectively and Lane 4 & Lane 5 contain RT-PCR yields of total-cell RNA using cDNA of *L. major* in both promastigote and amastigote forms respectively. Lane 3 contains a band near 255 bp; no detection of bands in Lane 2, Lane 4, and Lane 5. It was performed in the presence of a positive control (GAPDH, 496 bp) in Fig. 6.a.II. 6.b. Fluorescence microscopy did not detect ALSP in the promastigote (6.b.I) and amastigote (6.b.II) of *L. major* with antisera and FITC-conjugated anti-rabbit IgG secondary antibody at 40x magnification with mounting media containing DAPI and filters were used 461 nm for DAPI (stained nucleus) and 591 nm for FITC (stained ALSP).

It was also performed through fluorescence microscopy where the promastigote and amastigote of *L. major* were stained with the antisera and FITC-conjugated anti-rabbit IgG secondary antibody with mounting media containing DAPI. Fluorescence microscopy did not show expression of ALSP in either promastigote or amastigote of *L. major* (**Figure 6.b**).

### Purification and sequence analysis of ALSP from *Leishmania donovani* amastigotes

The antisera against ALSP was used to purify the native protein from the amastigotes’ lysate. Both the preimmune and antisera were cross-linked with protein A-Sepharose bead (Invitrogen-101041). The purification was performed with another negative control, albumin-like protein. Both the negative controls could not detect the protein. When probed, the lysate of amastigotes revealed a signal at 10 kDa (**Figure 7.a**). The precise molecular weight obtained by MALDI-TOF MS was 10.147 kDa (**Figure 7.b**). MALDI-TOF-MS-MS analysis showed the sequence with 100% identity with leishmanial ALSP (CBZ37742.1), using the peptide fragments of the trypsin-digested protein (from amastigotes) (**Figure 7.c**).

**Figure. 7.**
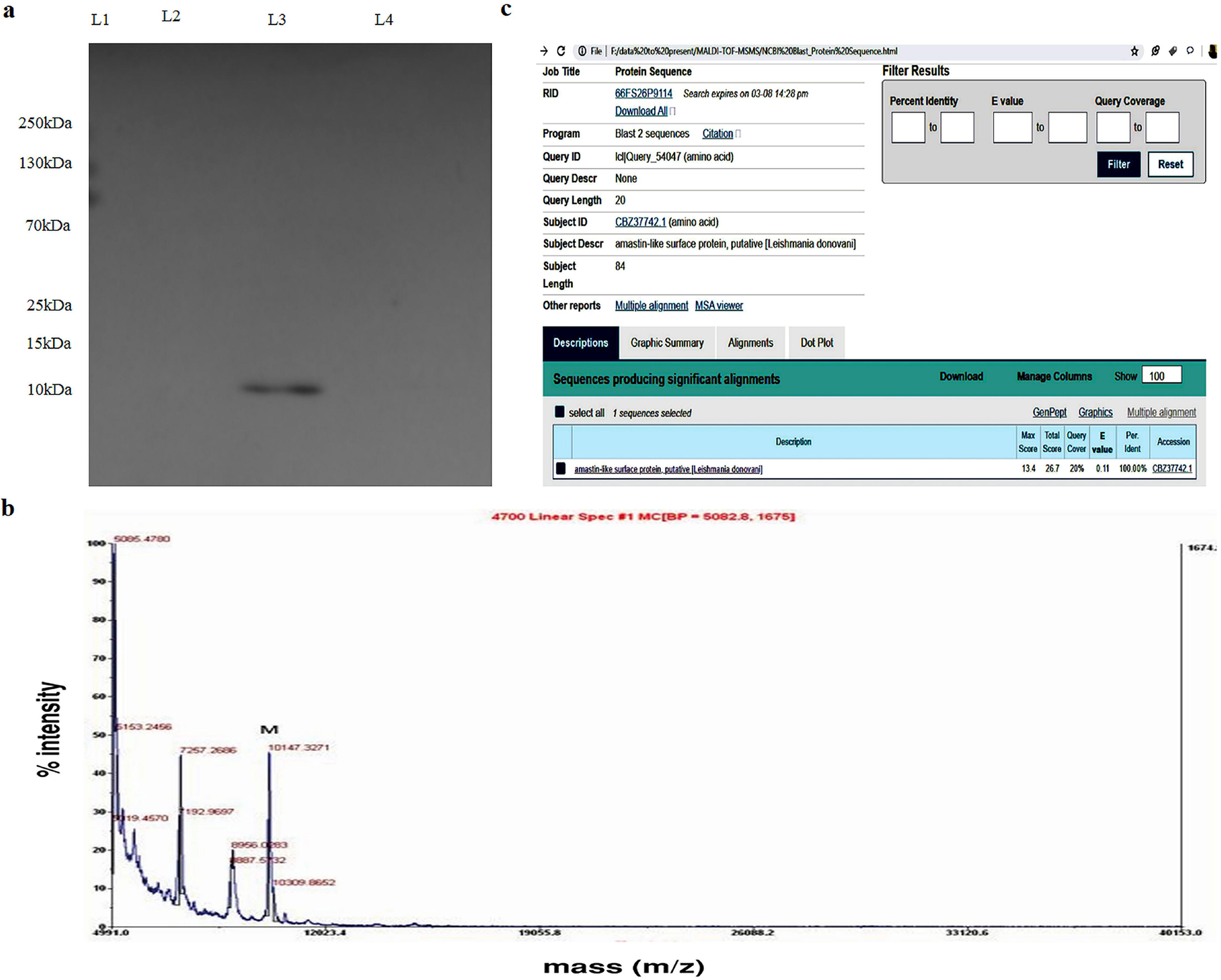
Western blot of ALSP expression and determination of molecular weight of ALSP by MALDI-TOF. (7.a) Western blotting was carried out with antiserum and anti-rabbit horseradish peroxidase-conjugated secondary antibody. Lane 1, molecular mass marker (10-250 kDa); Lane 2, purified protein expressed in promastigotes, used as negative control; Lane 3, ALSP band at 10 kDa in amastigotes; Lane 4, albumin-like protein expressed in amastigotes, used as negative control. (7.b) The M peak at 10.147 kDa revealed the accurate molecular mass of ALSP by MALDI mass spectrometry. (7.c) Sequencing of the purified protein was done by MALDI-TOF-MS-MS analysis after tryptic digestion. The sequencing data of ALSP after MALDI-TOF-MS-MS analysis showed 100% identity with leishmanial amastin-like surface protein (ALSP) when uploaded in the NCBI for BLASTn

## DISCUSSION

The results of the current study preliminary suggest that a novel protozoan parasitic protein, amastin-like surface protein (ALSP), acts as a lipase enzyme. The function of the protein was predicted through *in-silico* analysis by I-TASSER. The *in-vitro* analysis with lipase assay ravelled to produce fatty acid in amastigotes. Further, to characterize the protein, it was cloned in a bacterial system and a polyclonal antibody was produced against the 10 kDa protein, which was present in *L. donovani* amastigote. Immunofluorescence study showed that the expression of amastin-like surface protein in amastigotes but not in the promastigotes. The protein was unexpressed in *L. major*.

While evaluating the function of the target novel protein amastin-like surface protein (ALSP), the I-TASSER software highly predicted the role of the putative protein as a triacylglycerol lipase. Closer examination of the peptide sequence revealed the protein as serine-rich, where the signature of Class III serine lipases belonging to numerous *Leishmania* species. In our preliminary experiments, we could demonstrate the lipase activity of the amastin-like surface protein. This protein had the spaced trypsin-like catalytic triad, Ser-His-Asp ^[29]^. Closer examination of the peptide sequence also did not reveal the commonly described sequence of glycine-x1-serine-x2-glycine (x usually is histidine or tyrosine), the common catalytic sequence present in a wide variety of lipases. However, a GAS domain was present in ALSP. GHS domains were present in eleven other putative lipases of *Leishmania donovani* reported in the GenDB database (data not shown). +The amastin-like surface protein (ALSP) has serine richness and also have histidine and glycine distribution, as well as terminal serines characteristic of lipases.

Lipases are an important class of enzymes, contributing to the exoproteome of multiple *Leishmania* species. Earlier, such a lipase was examined in *Leishmania* major [Freidlin] [30]. In the current study, either the script or the mature ALSP protein in *L. major* was not detected. Our cross-examination of multiple lipases from *L. donovani* from the GenDB database did not show major overlap and sequence alignment with ALSP, likely supporting our claim that ALSP is a novel lipase of *L. donovani*. The GenDB database additionally shows lipases present in multiple other species of *Leishmania* and other kinetoplastids including multiple species of *Trypanosoma*. These lipases are also present in many fungi, as well as prokaryotic organisms like *Yersinia, Pseudomonas cepacia*, and *Helicobacter pylori* ^[31] [32] [33]^

What it means to be a functional lipase in *Leishmania* is far from clear but there is few precedence as to the role of the lipases. These are simple hydrolases generating glycerol and fatty acids, which are important substrates for metabolic processes. Furthermore, these are important membrane components and lysis of membrane may be one of the first major steps in insinuation of a deflagellated promastigote into the host phagolysosome to form an amastigote to accomplish its digenic lifecycle. This however may not be the role for ALSP, as ALSP expression was seen minimal to non-existent in the promastigote form in the present study. A previous preliminary study also could not detect any low molecular protein below 20 kDa in *L. donovani* promastigotes ^[34]^. Earlier though it was reported that the lipase *LdLip3* was present in both the promastigote and the amastigote forms of *L. major*^[30]^, where it might contribute to vertebrate host cell entry, as well as contribute to virulence mechanisms like membrane remodelling of the phagolysosome.

Parasites rely on a complex system of uptake and synthesis mechanisms to satisfy their lipid needs. Parasites like *Leishmania* have adopted complex mechanisms of host lipid harvesting ^[35]^. The glycerol from hydrolysed lipid is transported across to the parasite from the host cell through aquaporin AQP1-like aquaglyceroporin channels, contributing to the complex energy metabolism of the parasite ^[36]^. Glycerol-transport proteins have been predicted to be present in *L. donovani* ^[37]^. Fatty acids, derived from the host, are transported back to the parasite by complex mechanisms ^[18] [38]^. Apart from supporting parasite energy generation, the fatty acids may be importantly involved in the neo synthesis of special membrane lipids like phosphorylceramide ^[15]^. Recently, a preliminary study has demonstrated a role for albumin-like protein in the uptake of fatty acids by the intracellular form of *L. donovani*^[27]^ Importantly, these channel proteins are significantly targeted by antimonials, one of the most important chemotherapeutic avenues for treating advanced leishmanial diseases. Though chemotherapy options like antimonials, amphotericin B, and miltefosine remain the mainstay for treating visceral leishmaniasis, emerging drug resistance, non-availability of drugs, and toxicity remain major obstacles in obtaining complete remission and significantly contributes to sustained mortality of visceral leishmaniasis. Thus, attempts for developing vaccines have been aimed.

ALSP does not have a KDEL sequence, making it unlikely to dock to a cell membrane or an organelle membrane and shall function as a lipase. Normally, lipases are relatively bulky proteins with an approximate molecular weight of 60 kDa ^[39]^. The predicted molecular weight of the putative lipases of *L. donovani* were around 70 kDa (As seen in UniProt). *LdLip3* molecular weight is 33 kDa. Recombinant *Leishmania* antigens (single peptides/polypeptides) was used to produce second-generation vaccines. Among different trials, promising results in phase I were shown by GLA-SE adjuvant tagged LEISH-F3, a multicomponent vaccine, to produce an immune response in healthy subjects ^[40]^. The molecular weight determination of ALSP revealed a rather relatively small protein of 10 kDa. This makes ALSP an important target as a peptide vaccine component. Preliminary examination with VaxiJen v2.0 and ANTIGENPro shows the immunoinformatic feasibility of eliciting cell-based immunity against this peptide target. Our cloning studies revealed the potential of generating the ALSP protein in large quantities *in-vitro*. This remains our goal for future studies. Cumulatively, the results of the present study show that the novel 10 kDa ALSP supports *Leishmania donovani* parasitism by its lipolytic activity in the amastigote form.

As *Leishmania* is an obligate parasite, it must procure macromolecules like glycerol and fatty acids to facilitate their opportunistic survival. Previous studies have shown the preference of amastigotes in using fatty acids as their carbon source via beta-oxidation, apart from utilizing glucose and proline ^[35] [38] [41] [42]^. *LdLip3* was shown to be active in both promastigotes and amastigotes of *L. donovani* ^[30]^ whereas ALSP activity was negligible in promastigotes of *L. donovani*. ALSP may facilitate the survival of the parasite by remodelling of the phagolysosome membrane and may induce tissue inflammation when secreted in the human host. Earlier studies have shown that lipase precursor-like protein confers the oral drug alkylphosphocholine miltefosine (but not sodium antimony gluconate and amphotericin B) resistance in *L. donovani* by enhancing macrophage infectivity and increasing IL-10/TNFα ratio ^[43]^. The solitary expression of ALSP in amastigotes rather than promastigotes is an exceptional phenomenon, as only a percentage of proteins are differentially expressed between these morphological forms. For the most part, all proteins are constitutively expressed in both promastigotes and amastigotes, making it non-selective for the parasite to survive in the invertebrate and human host ^[1] [37]^. Additionally, bioinformatics prediction and the current study indicates differential expression of ALSP in different species of *Leishmania*(*donovani vs. major*), which also is an intriguing observation.

Surprisingly, there is a significant overlap between diverse lipases and esterase enzymes acetylcholine esterase carboxylesterase, and cholesterol esterase ^[39]^. Though the geometry of the active size varies, as also seen with ALSP, there is overlap between these classes of enzymes. The nomenclature of amastin-like surface protein (ALSP) does not suggest a similarity with amastin proteins, a significant class of surface proteins widely expressed in all leishmanial species, *Trypanosoma brucei*, responsible for sleeping sickness and *Trypanosoma cruzi*, the inducer of Chagas disease. Preliminary sequence alignment did not show overlap with alpha, beta, gamma, or delta amastins (data not shown). Though amastin proteins are widely distributed, detailed investigations into the function of amastins are only scanty. Secondary structure comparison of amastins has shown its similarity to claudins, important cell-cell junction proteins ^[44]^. Preliminary reports suggest that the amastigote form of *T. cruzi* develops cell synapses before the insinuation in the secondary host cell ^[45]^. Whether ALSP performs such roles remains unresolved in the current study. Intriguingly, lipases are related to some cell adhesion proteins like *Drosophila* neurotactin ^[46]^.

Advanced leishmanial disease often present with bleeding diathesis, making it difficult to obtain tissue biopsies for establishing diagnosis. Examination of ALSP in the peripheral blood may aid as a diagnostic tool under such circumstances. Though initial vaccination strategies have not received robust outcomes, it may be appreciated that efforts to develop new vaccines are critical. HIV-leishmanial coinfections are important emerging infections across the globe in both rural and urban areas. These important theragnostic implications elevate the significance of further examining the novel protein ALSP in future studies.

## AUTHOR CONTRIBUTIONS

BB, Performed majority of experiments and drafted first version of manuscript; BL, coordinated MALDI-TOF experiments; PS, Insilico studies, and Data Analysis, KD, Draft preparation, MG, Conceptualized the study, obtained funding, supervised experiments, coordinated drafting of manuscript.

## ACKNOWLEDGMENT

A preliminary version of the manuscript was submitted in the BioRxiv repository (https://doi.org/10.1101/2020.07.23.218107).

